# Large-scale Analyses of Disease Biomarkers and Apremilast Pharmacodynamic Effects

**DOI:** 10.1101/652875

**Authors:** Irina V. Medvedeva, Matthew E. Stokes, Dominic Eisinger, Samuel T. LaBrie, Jing Ai, Matthew W. B. Trotter, Peter Schafer, Robert Yang

**Affiliations:** Celgene Corporation, Informatics&Predictive Sciences, Cambridge, 02140, USA; Myriad RBM Inc., Austin, 78759, USA; Celgene Corporation, Celgene Institute for Translational Research Europe (CITRE), Sevilla, 41092, Spain; Celgene Corporation, Translational Development, Summit, 07901, USA

## Abstract

Finding biomarkers that provide shared link between disease severity, drug-induced pharmacodynamic effects and response status in human trials can provide number of values for patient benefits: elucidating current therapeutic mechanism-of-action, and, back-translating to fast-track development of next-generation therapeutics. Both opportunities are predicated on proactive generation of human molecular profiles that capture longitudinal trajectories before and after pharmacological intervention. Here, we present the largest plasma proteomic biomarker dataset available to-date and the corresponding analyses from placebo-controlled Phase III clinical trials of the phosphodiesterase type 4 inhibitor apremilast in psoriasis (PSOR), psoriatic arthritis (PsA), and ankylosing spondylitis (AS) from 526 subjects overall. Using approximately 150 plasma analytes tracked across three time points, we identified IL-17A and KLK-7 as biomarkers for disease severity and apremilast pharmacodynamic effect in psoriasis patients. Combined decline rate of KLK-7, PEDF, MDC and ANGPTL4 by Week 16 represented biomarkers for the responder subgroup, shedding insights into therapeutic mechanisms. In ankylosing spondylitis patients, IL-6 and LRG-1 were identified as biomarkers with concordance to disease severity. Apremilast-induced LRG-1 increase was consistent with the overall lack of efficacy in ankylosing spondylitis. Taken together, these findings expanded the mechanistic knowledge base of apremilast and provided translational foundations to accelerate future efforts including compound differentiation, combination, and repurposing.

## Introduction

Psoriasis (PSOR) is an autoimmune disease and manifests as thickened erythematous patches of skin covered with silvery scales that can arise across the entire body but often at predisposed areas such as the extensor aspects of elbows and knees, nails, scalp, palms, soles, and intertriginous areas. The psoriatic lesions are frequently associated with pain and pruritus^1, 2^. Histologically, the skin lesions are characterized by acanthosis from rapid keratinocyte proliferation, hypogranulosis, parakeratosis from aberrant keratinocyte differentiation, erythema caused by dilation of blood vessels in the papillary dermis, and a dense inflammatory infiltrate consisting of dendritic cells and T cells and dendritic cells. Psoriatic arthritis (PsA) is a chronic inflammatory arthritis present in up to 42% of individuals with PSOR, and is classified as one of the seronegative spondyloarthropathies, based on the presence of spinal involvement in about 50% of patients^3^. In the majority of patients, PSOR precedes PsA by several years, especially in patients with nail involvement^4^. PsA is manifested as one of 5 subtypes: distal interphalangeal joint involvement only, asymmetrical oligoarthritis, symmetrical polyarthritis, spondylitis, and arthritis mutilans^5^. Ankylosing spondylitis (AS) is a systemic chronic, inflammatory disorder primarily affecting the spine and characterized by sacroiliitis and lower back pain that can result in destruction and fusion of spinal vertebrae^6^. PSOR, PsA, and AS are pathologically related conditions sharing common clinical manifestations and immunological drivers characterized by dendritic cells and helper T cells producing Th1 and Th17 cytokines including IL-23, TNF-α, IFN-γ, IL-17, and IL-22^7–9^.

Apremilast, a selective small-molecule inhibitor of phosphodiesterase 4 (PDE4), is an approved oral drug for psoriasis and psoriatic arthritis^10^. In previous studies it was shown that apremilast reduces circulating levels of Th1 and Th17 proinflammatory mediators and increases anti-inflammatory mediators in these diseases^11^. Phosphodiesterase 4 is a cyclic adenosine monophosphate (cAMP)-specific PDE and the dominant PDE in inflammatory cells. Inhibition of PDE4 elevates intracellular cAMP levels, which in turn downregulates the inflammatory response by modulating the expression of TNF-α, IL-23, IL-17, and other proinflammatory cytokines. Elevation of cAMP also increases anti-inflammatory cytokines including IL-10^12^. Consistent with these causal mechanisms, TNF-α and IL-17A was known to be upregulated in PSOR and downregulated by apremilast^13^. In an open-label phase II PSOR study, apremilast decreased lesional skin epidermal thickness and expression of pro-inflammatory genes, including inducible nitric oxide synthase (iNOS), IL-12/IL-23 subunit p40, IL-23 subunit p19, IL-17A, IL-22, and IL-8^10^. In a placebo-controlled double-blinded phase III PSOR trial (ESTEEM2), treatment with apremilast 30 mg BID resulted in significant reductions of plasma IL-17A, IL-17F, IL-22, and TNF-α protein levels^14, 15^. In a placebo-controlled double-blinded phase III study in PsA, apremilast treatment was associated with decreased plasma levels of IL-6, IL-8, MCP-1, MIP-1β, TNF-α, ferritin, and a small increase in von Willebrand factor (vWF) plasma protein levels. Among these, the changes in TNF-α and vWF were significantly associated with achieving an American College of Rheumatology 20 (ACR20) clinical response. After 40 weeks of apremilast treatment, significant decreases in plasma IL-6, IL-23, IL-17, and ferritin, and significant increases in IL-10 and IL-1RA were observed^11^.

Taken together, these results suggest that partial inhibition of several proinflammatory mediators, including TNF-α and IL-17 may play a key role in the mechanism of action of apremilast in the treatment of psoriatic diseases. Expansion of larger protein biomarker sets that link disease and drug effect will help organizing next-generation effective therapies.

## Results

The abundance of 155 plasma proteins including chemokines and cytokines was measured by Myriad RBM sandwich immunoassays and analyzed in ankylosing spondylitis, psoriatic arthritis, or psoriasis subjects who participated in 3 independent Phase III apremilast clinical trials (POSTURE, ESTEEM2, PALACE1). Briefly, subjects were randomly selected to have received apremilast (20mg or 30 mg) or placebo during the first 16 weeks of respective trials, and assessed for response status according to pre-specified end points. For all post-hoc analyses in the present study, 20 mg and 30 mg arms were combined together (Table 1). Myriad RBM data were collected from subject plasma upon each visit at baseline, Week 4 and Week 16 (details in Methods).

**Table 1.**
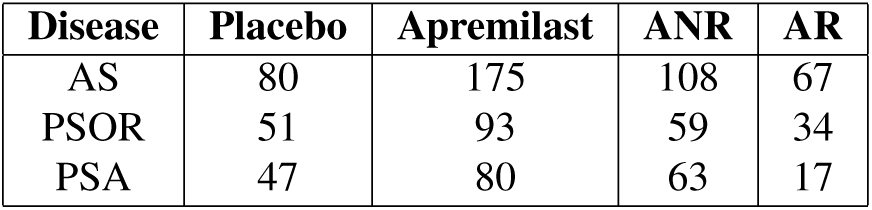
Number of the patients enrolled in the study in each arm and participated in protein production data collection. ANR - apremilast non-responders; AR - apremilast responders.

| Disease | Placebo | Apremilast | ANR | AR |
| --- | --- | --- | --- | --- |
| AS | 80 | 175 | 108 | 67 |
| PSOR | 51 | 93 | 59 | 34 |
| PSA | 47 | 80 | 63 | 17 |

### Systematic discovery of disease biomarkers

We set out to determine the most prominent plasma protein biomarkers of disease severity, for each disease. Lasso regression with L1 regularization was applied to select proteins with the largest independent correlation coefficient to severity. The analyses were carried out initially at baseline, and repeated for later time points to estimate the robustness. From a panel of 122 plasma proteins that passed QC, 10 showed various degrees of correlation with PASI scores within the psoriasis subjects. When cross-referenced with later time points where PASI scores changed for some individuals, KLK-7 and IL-17A remained as the most consistent and robust correlates based on effect size and directionality (Figure 1A). While IL-17A was a well known marker, KLK-7 represented a novel psoriasis plasma biomarker. Similar analyses were performed for ankylosing spondylitis and psoriatic arthritis subjects. LRG-1 was best associated with ASDAS score in patients with ankylosing spondylitis (Supplement Figure 1). No consistent plasma proteins were found to be correlated with DAS28 score in the psoriatic arthritis cohort.

**Figure 1.**
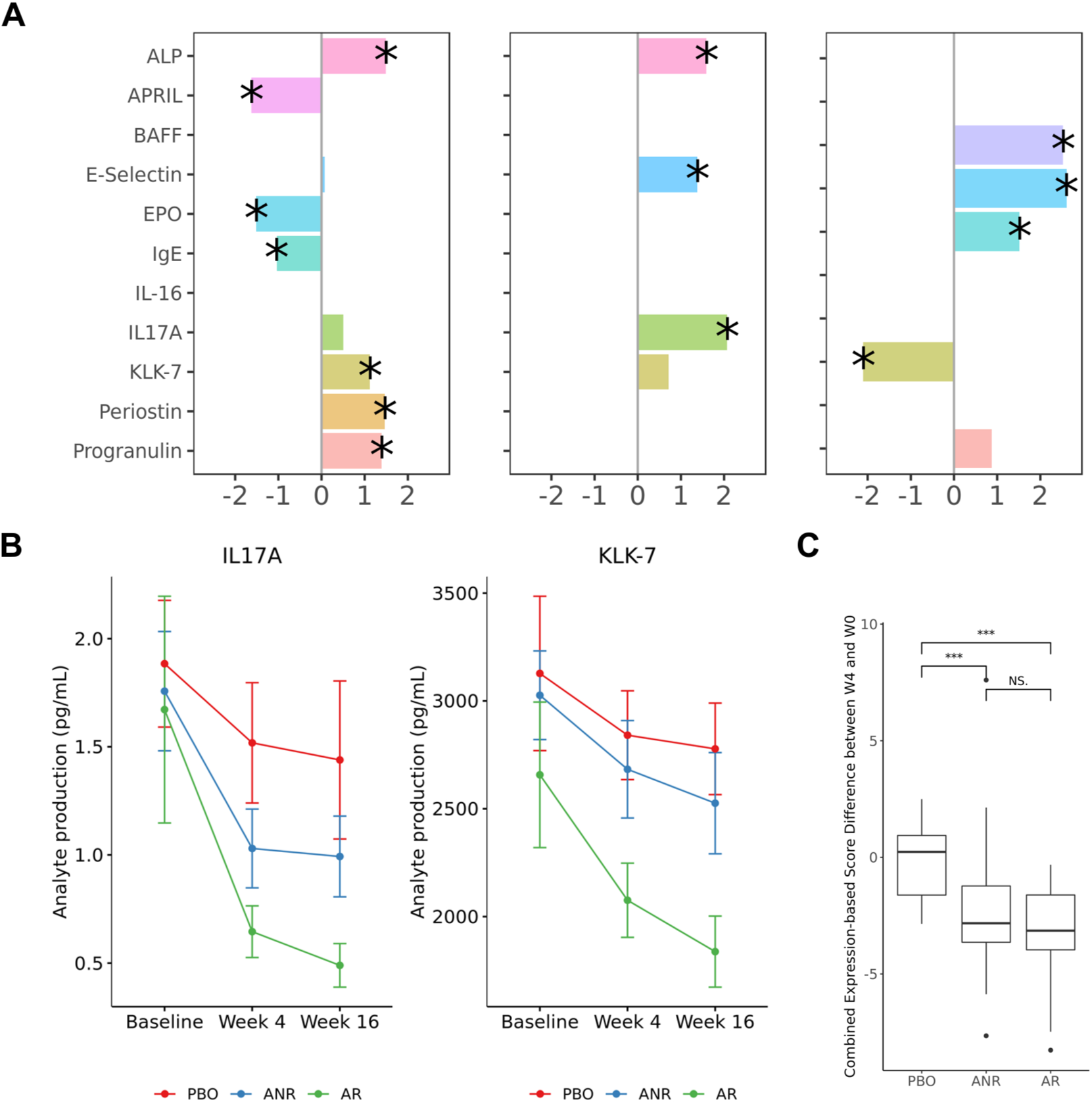
IL-17A and KLK-7 as biomarkers of psoriasis. A) Lasso coefficients of analytes to PASI score. Only proteins that were significant in at least one time point are shown. Stars indicate significance (p-value < 0.05). B) Mean analytes concentration of IL-17A and KLK-7 in pg/mL for each subgroup across time points; C) Comparison of the change in aggregated IL-17A and KLK-7 expression score in the first 4 weeks in subgroups: placebo (PBO), apremilast responders (AR) and apremilast non-responders (ANR) (*** denotes p<0.01). Aggregated score was significantly lowered by apremilast.

Following the initial exploration, we selected prominent analytes to understand the relationships of IL-17A and KLK-7 in psoriasis and LRG-1 in ankylosing spondylitis subjects, respectively. In the psoriasis cohort, we confirmed the correlations to PASI scores were independent from subject demographics by univariate regression analyses controlling for age and gender (Table 2). The quantitative change in PASI scores tracked well with the magnitude of change in analyte expressions, referred to as the delta-delta correlations (Supplementary Tables 1-3). Furthermore, the pharmacodynamic effect of both analytes differ significantly between the placebo, apremilast responder and non-responder groups (Figure 1B). Similarly in the ankylosing spondylitis cohort, LRG-1 was confirmed to be significantly correlated with ASDAS score (Table 3), and in delta-delta analyses (Supplementary Tables 4-6), independent of age and gender.

**Table 2.** Beta coefficients, p-values and *R*^2^ values in linear regression models evaluating correlation between PASI total score and protein expression in PSOR patients.

| PASI<br>Analyte | Baseline |  | Week 4 |  | Week 16 |  |
| --- | --- | --- | --- | --- | --- | --- |
|  | beta | p | beta | p | beta | p |
| *IL-17A* | 3.46 | 2.73E-08 | 3.85 | 4.81E-07 | 4.85 | 4.01E-08 |
| gender | 0.98 | 0.3715 | 2.30 | 0.0705 | 1.57 | 0.29 |
| age | -0.06 | 0.3282 | 0.06 | 0.9929 | 0.07 | 0.9933 |
| $R^2$ | 0.29 | | 0.32 | | 0.35 | |
| p(F-test) | 2.47E-07 |  | 1.20E-06 |  | 1.54E-07 |  |
| *KLK-7* | 3.36 | 1.64E-07 | 3.52 | 1.56E-05 | 5.04 | 6.32E-09 |
| gender | 0.18 | 0.88 | 1.55 | 0.2452 | 1.38 | 0.34 |
| age | -0.02 | 0.5764 | 0.06 | 0.9929 | 0.05 | 0.9933 |
| $R^2$ | 0.26 | | 0.26 | | 0.38 | |
| p(F-test) | 1.34E-06 |  | 3.21E-05 |  | 2.65E-08 |  |
| *Additive IL17A+KLK7* |  |  |  |  |  |  |
| IL17A | 2.40 | 3.97E-05 | 1.63 | 7.91E-3 | 1.26 | 3.73E-2 |
| KLK7 | 2.06 | 5.11E-4 | 1.99 | 1.64E-3 | 1.85 | 2.31E-3 |
| gender | 0.31 | 0.76 | 0.06 | 0.96 | -0.47 | 0.67 |
| age | -0.04 | 0.3264 | 0.01 | 0.7399 | 0.01 | 0.77 |
| $R^2$ | 0.35 | | 0.26 | | 0.20 | |
| p(F-test) | 2.63E-11 |  | 1.36E-06 |  | 4.75E-05 |  |

**Table 3.** Beta coefficients, p-values and *R*^2^ values in linear regression models evaluating correlation between ASDAS score and protein production in AS patients.

| ASDAS<br>Analyte | Baseline |  | Week 4 |  | Week 16 |  |
| --- | --- | --- | --- | --- | --- | --- |
|  | beta | p | beta | p | beta | p |
| *Univariate IL-6* | 0.43 | 3.66E-18 | 0.45 | 5.13E-14 | 0.48 | 1.45E-13 |
| gender | 0.05 | 0.5895 | -0.09 | 0.6804 | -0.07 | 0.6160 |
| age | -0.10 | 0.1598 | -0.16 | 0.0921 | -0.08 | 0.9066 |
| $R^2$ | 0.30 | | 0.25 | | 0.23 | |
| p(F-test) | 3.49E-18 |  | 2.64E-13 |  | 4.25E-12 |  |
| *Univariate LRG1* | 0.53 | 2.56E-32 | 0.47 | 1.14E-16 | 0.5 | 4.15E-15 |
| gender | 0.27 | 0.0575 | 0.14 | 0.6804 | 0.18 | 0.5291 |
| age | -0.13 | 0.1239 | -0.17 | 0.0921 | -0.03 | 0.9066 |
| $R^2$ | 0.46 | | 0.29 | | 0.26 | |
| p(F-test) | 3.79E-32 |  | 6.81E-16 |  | 1.35E-13 |  |
| *Additive IL6+LRG1* |  |  |  |  |  |  |
| IL6 | 0.14 | 2.75E-03 | 0.24 | 2.52E-04 | 0.26 | 3.65E-04 |
| LRG1 | 0.44 | 1.84E-18 | 0.32 | 2.30E-07 | 0.33 | 4.50E-06 |
| gender | 0.21 | 0.0143 | 0.03 | 0.8136 | 0.05 | 0.6603 |
| age | -0.12 | 0.0432 | -0.16 | 0.0373 | -0.05 | 0.51 |
| $R^2$ | 0.49 | | 0.33 | | 0.29 | |
| p(F-test) | 1.45E-38 |  | 4.88E-20 |  | 1.28E-17 |  |

Next, we explored the possibility of additive effects by evaluating the strength of combining individually biomarkers. We showed that the additive model of IL-17A and KLK-7 significantly captured more PASI score variability than univariate models in psoriasis (Figure 2, Supplementary Table 7). In an attempt to link the combined disease biomarkers with potential clinical utilities, we constructed a simple aggregated biomarker score to monitor apremilast effect. The values of coefficients from the baseline were used as weights to calculate the combined score. In psoriasis subjects, the combined score of IL-17A and KLK-7 was significantly downregulated after apremilast treatment (Figure 1C). By contrast, in AS subjects, we included IL-6 along with LRG-1 based on their significant univariate correlations to ASDAS score. However, the IL-6 + LRG1 additive model showed a strong association that was not significantly stronger than LRG-1 alone (Supplementary Tables 8). This finding suggested that IL-6 correlation was likely related to that of LRG-1, consistent with the Lasso feature selection analysis where IL-6 was not selected in the presence of LRG-1. Despite the high association with ASDAS disease scores, the combined IL-6 + LRG-1 score in ankylosing spondylitis showed little apremilast-induced pharmacodynamic effect. The link with disease but not apremilast effect may offer a plausible molecular connection consistent with an overall lack of apremilast clinical response in this trial (Supplementary Figure 2).

**Figure 2.**
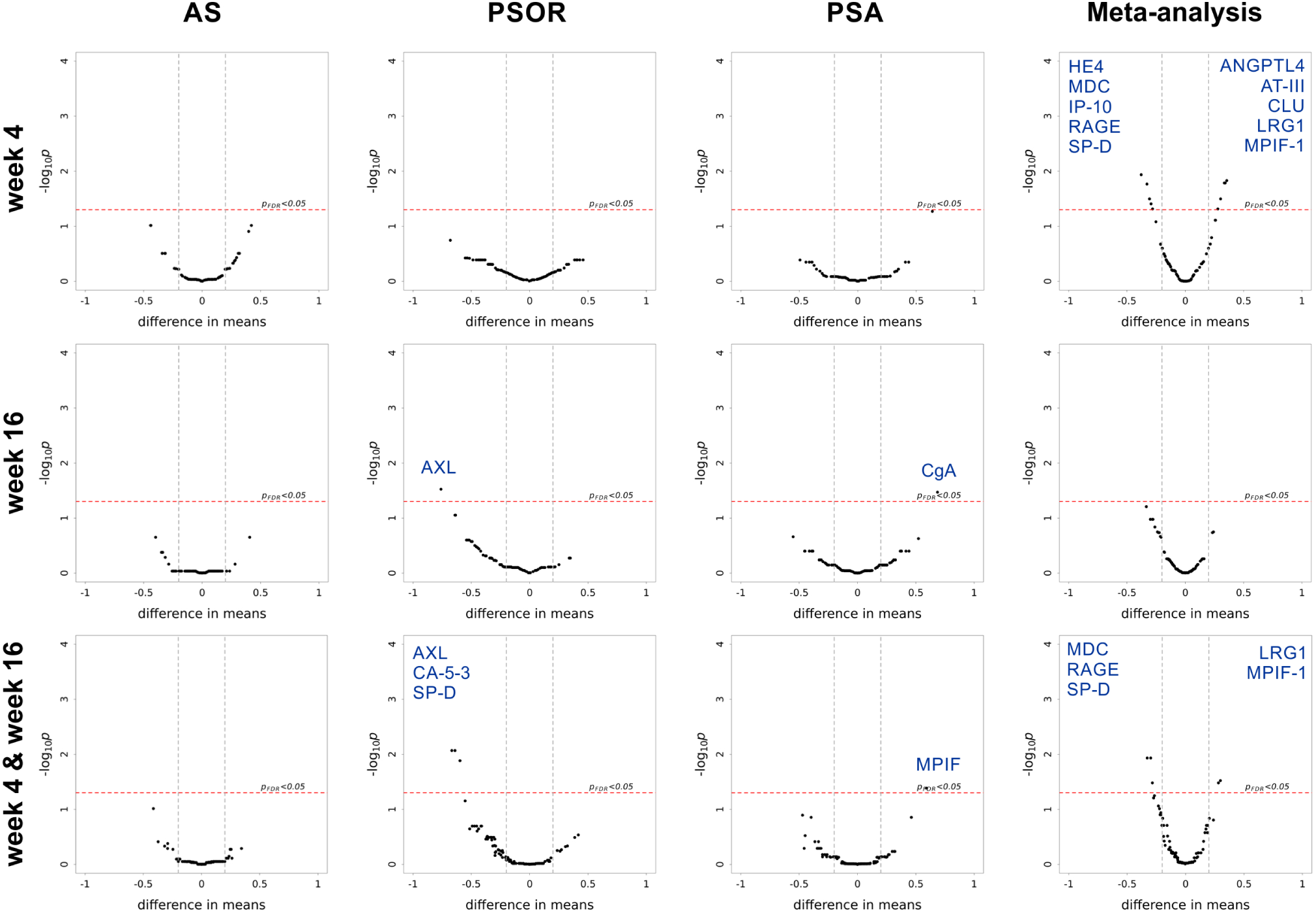
Differential expression (DE) analysis between placebo and treatment arms at specified time points and meta-analysis across all diseases. Significant DE proteins are labeled. The last row “week4 & week16” denotes to mixed-effect modeling on pooled data from the two time points. The last column “Meta-analysis” denotes to differential expression modeling on pooled data across all three diseases.

### Apremilast effects at each time point

We next examined the temporal and overall biomarker patterns corresponded to the apremilast pharmacodynamic effect and response differences. Differential expression analyses showed very few analytes that were robustly different between placebo and apremilast when comparing at weeks 4 and 16 separately (Figure 2). Pooling together both time points and modeling by mixed-effect models increased the detection power within each disease. In psoriasis subjects, AXL, CA-15-3 and SP-D were significantly downregulated by apremilast. Of note, AXL and its upstream stimulus, IL-16 are known to be involved in Th1^16, 17^, Treg^18^, and Th17^19^ regulation. In psoriatic arthritis subjects, MPIF-1 (CCL23) was upregulated in the apremilast arm. MPIF-1 is a chemokine that induce epithelium cell migration while reducing the proliferation of myeloid progenitor cells^20^, and is produced by neutrophils in response to TLR-agonists and TNFα expression^21^.

These individual subgroup comparisons of placebo and apremilast arms per disease per time point did not converge on a common set of apremilast pharmacodynamic protein markers across diseases. We reasoned that meta-analyses combining diseases and timepoints may provide additional detection power. As shown in Figure 2, meta-analysis combining 3 diseases at Week 4 revealed a set of temporal pharmacodynamic biomarkers implicated in autoimmune implications and known to impact inflammation states within the Th17/Th1 axis. Specifically, HE4, MDC, IP-10, RAGE and SP-D were significantly down-regulated by apremilast, while ANGPTL4, AT-III, CLU, LRG-1 and MPIF-1 were upregulated. MDC was recently reported among biomarkers of atopic dermatitis severity^22^, a disease with overlapping clinical symptoms as psoriasis. RAGE (S100A12), a Th17 pathway-dependent protein produced by keratinocytes^23^, was reported to be upregulated with β2m-amyloidosis in patients with rheumatoid arthritis^24^. Surfuctant protein D is known to be upregulated in lesional psoriatic skin25. IP-10, a known biomarker of localized scleroderma26, psoriasis and psoriatic arthritis27, is produced mostly by Th1 cells28. In contrast, meta-analyses combining across diseases for Week 16 did not yield significant biomarkers, indicative of larger heterogeneity at a longer time frame that may be confounded due to a combination of pharmacodynamic and disease recovery response to treatment. To determine pharmacodynamic markers with sustained signals (combine weeks 4 and 16) at higher detection power, we constructed mixed-effect model on the pooled data across diseases. Several analytes remained as sustained effect in the combined time points. Notably, LRG-1, positively correlated with worse severity in ankylosing spondylitis, was higher in the apremilast arm at Week 4, providing biomarker consistency with lack of clinical efficacy.

### Apremilast pharmacodynamic effect by time-series analyses

Differential analyses in the previous section elucidated the most robust relative differences between groups at each time snapshots (individual or pooled). Questions remain about overall trend directionality and rate of change. We therefore sought to capture the full dynamic patterns via time-series analyses. CoGAPS algorithm^29^, a method based on Markov chain Monte Carlo non-negative matrix factorization (MCMC NMF), was adapted to characterize transition patterns. When jointly applied with PatternMarkers^30^, analytes were probabilistically assigned to each pattern by calculating trajectory distance (see Methods).

Time-series analyses using data pooled across diseases resulted in 3 apremilast pharmacodynamic patterns (Figure 3A-B). LRG-1 was associated with the red pattern that was characterized by a differential Week 0 to 4 transition between the placebo and apremilast arms. Specifically, LRG-1 declined in the placebo group during the first 4 Weeks while apremilast induced an increase, further highlighting that the apremilast-induced LRG-1 effect may be even more undesirable over placebo given the biomarker moved towards the direction of marking for worse disease severity. In contrast, the other two CoGAPS patterns (green and blue) captured a “bump” or upregulation of analytes in the placebo group during the first 4 Weeks, while apremilast induced a sustained downregulation at through 16 weeks. Overall, 37 proteins were grouped in the blue pattern while uPar was the only one in the green pattern. Of note, IL-17A and RAGE of the blue pattern displayed the most robust signals, consistently associated with the same pattern in separate analyses for each individual disease (Figure 3C-D red pattern for the psoriasis trial, Supplementary Figure 5 A-B green pattern for the ankylosing spondylitis and C-D blue pattern for psoriatic arthritis trial).

**Figure 3.**
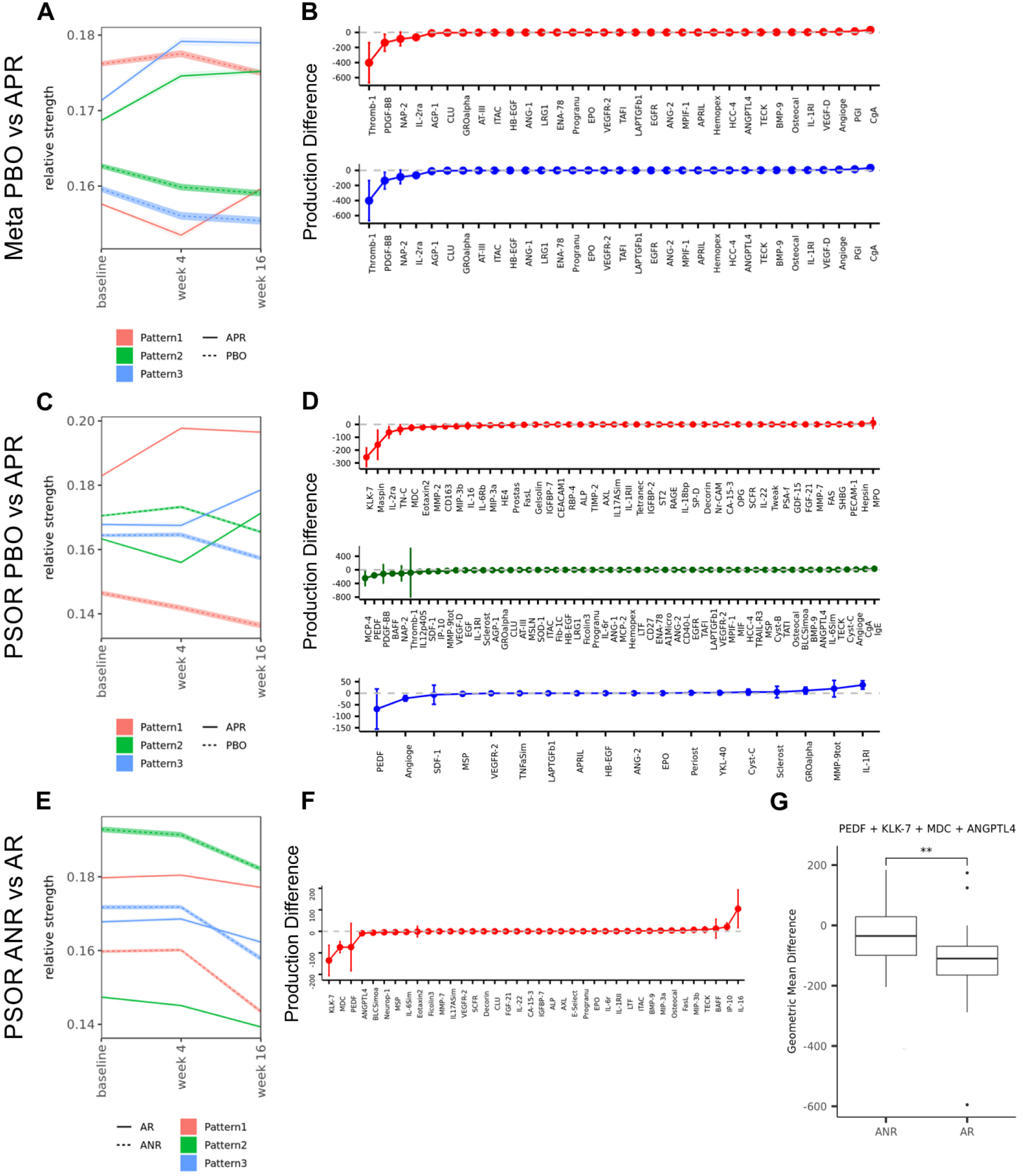
CoGAPS patterns across comparisons (A, C, F) and the associated analytes (B, D, G). The color of analytes and patterns is matched in each study. PSOR-psoriasis cohort; PBO - placebo treatment arm; APR - apremilast treatment arm; ANR - apremilast non-responders; AR - apremilast responders; Meta - meta-analysis on pooled data from all three diseases: ankylosing spondylitis, psoriasis and psoriatic arthritis. G) Geometric mean declines of PEDF, KLK-7, MDC, and ANGPTL4 (weeks 4 and 16) between apremilast non-responders and responders (** denotes p<0.05).

Similarly, we applied time-series analyses to compare apremilast response groups within the apremilast arms from each trial cohort. Several patterns emerged from the apremilast non-responder vs. responder comparison in the psoriasis trial (Figure 3E-F). One pattern (red) described a group of analytes that had subtle changes within the first 4 weeks, down-regulation by Week 16 regardless of clinical response, but seemingly with different rate of decline between the non-responder and responder groups. Interestingly, IL-17A and KLK-7 were among the analytes in this pattern. Using the original protein concentrations of these two biomarkers of interest, We confirmed the overall pattern (Figure 1B) and the characteristic that the transition rate in the first 4 weeks could not differentiate the response groups (Figure 1C). By themselves, the combined rate of decline of IL-17A and KLK-7 from Week 4 to 16 also did not significantly differentiate the response groups, which we reasoned was due to magnitude and insufficient representations. Subsequently, we selected the top 4 analytes ranked by the magnitude of mean declines in the apremilast arm from this pattern (PEDF, KLK-7, MDC, ANGPTL4, Figure 3F). Aggregated expression scores were calculated for each response group, and formally evaluated by Mann-Whitney U test (Figure 3G). We were able to confirm that the aggregated change from Week 4 to 16 represented quantifiable molecular concordance to clinical response. Application of the workflow to the other disease cohorts yielded no response-associated patterns.

## Discussion

Disease biomarker discoveries coupled with drug-induced pharmacodynamics studies are critical for improving patient benefits. The utilities of such linked research studies are two folds: patient selection for precision medicine and back-translation into next-generation of therapeutics. Data generation and subsequent analyses require joint efforts from pharmaceutical companies and technology partners in well-powered clinical trial settings. In this study we demonstrate the values of linking disease-centric and drug-centric biomarker studies through meta-analyses in 3 apremilast Phase III trials of psoriasis, ankylosing spondylitis, and psoriatic arthritis. Plasma analytes were profiled in pre- and post-treated participating subjects, and computational models were applied to explore implications to disease severity, apremilast effect and clinical response status.

In psoriasis subjects, we identified IL-17A and KLK-7 as robust biomarkers to PASI total severity scores, which were significantly downregulated by apremilast, making the two biomarkers uniquely positioned to link disease- and apremilast-centric biology. Furthermore, KLK-7 combined with 3 other analytes were declined significantly more in the responder group by Week 16. IL-17A had been previously reported as a known driver for psoriasis and its downregulation by apremilast was described as a molecular mechanism of action to explain efficacy. The present study confirmed these findings and added evidence that inadequate downregulation of IL-17A alone is insufficient to explain clinical non-response. KLK-7, a serine protease involved in skin barrier functions, displayed a similar disease- and pharmacodynamic-relevance as IL-17A, prompting the consideration of its causal role in psoriasis manifestation and apremilast efficacy. A systematic literature survey revealed a general association to skin-related diseases (Figure 4), including key publications on its dysregulation in atopic dermatitis^31, 32^ and psoriasis^33^. To our knowledge, the current profiling effort is the largest study to confirm misregulation of KLK-7 in psoriasis subjects, and the first report of linkage to the apremilast-induced effect. For broader validation, we surveyed KLK-7 mRNA expression patterns in human psoriasis subjects across public gene expression datasets with and without treatment (Figure 5). Upregulation of KLK-7 mRNA in peripheral and lesional skin was observed across multiple studies, consistent with the observed proteomic findings in the present psoriasis cohort. Interestingly, two other approved therapeutics for psoriasis, broadalumab (anti-IL-17A) and etanercept (anti-TNF), also downregulate KLK-7 mirroring apremilast-induced effect, thus providing plausible explanations of converged therapeutic mechanism. Taken together, these findings support KLK-7 as a therapeutic biomarker with a potential causal mechanism for disease etiology. As an example for back-translation utilities, we propose to monitor IL-17A and KLK-7 protein levels as preclinical surrogates to estimate the effectiveness of next-generation psoriasis therapy.

**Figure 4.**
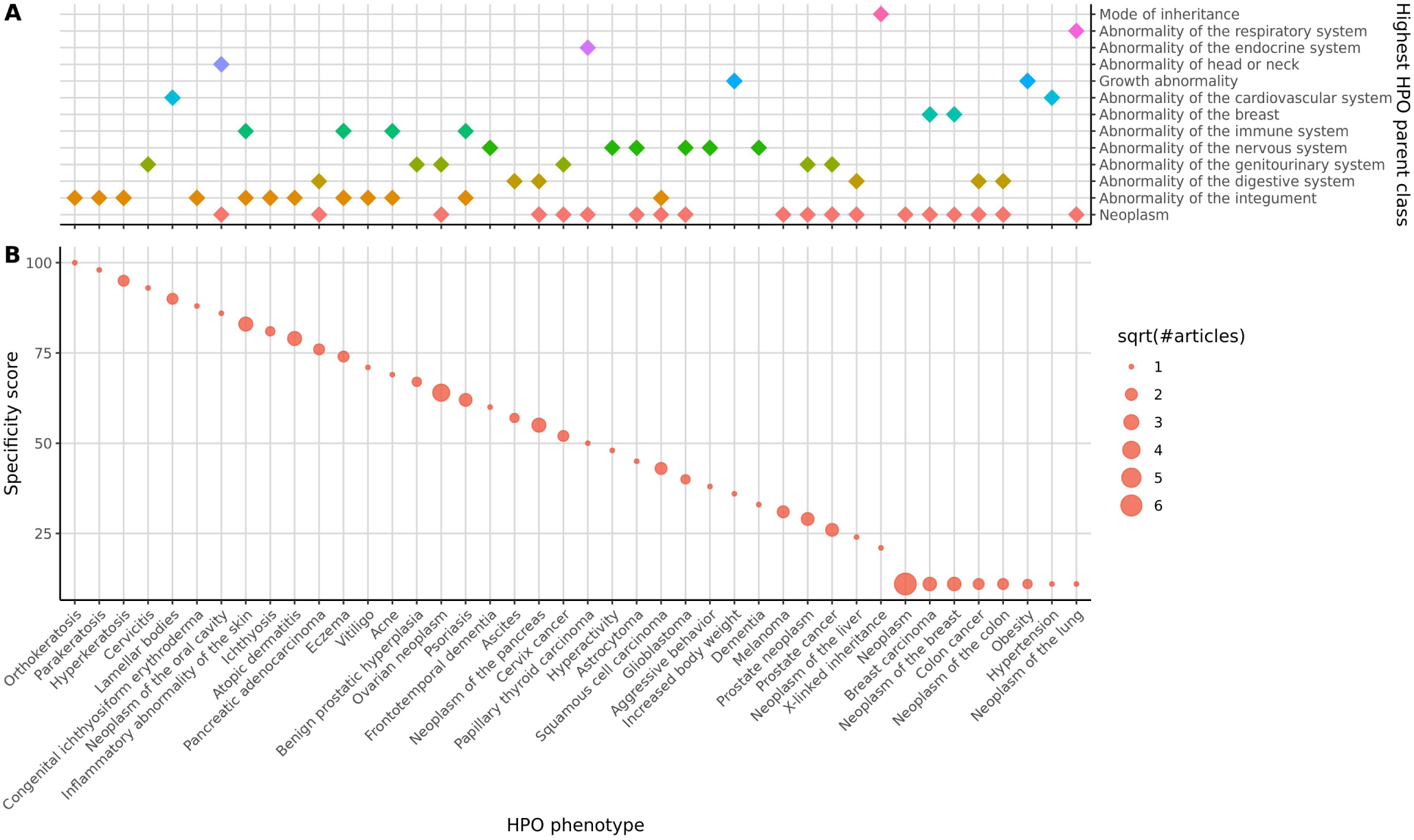
Systematic literature review of KLK-7 and phenotypes. Columns of panel A and B are aligned and labeled in panel B. A) The Highest HPO parent class(es) for each phenotype that co-occurred with KLK-7 in Medline abstracts. The association frequency of KLK-7 with the abnormality of the integument is the second highest behind neoplasm; B) Specificity score for each unique gene-phenotype pairing extracted from Medline abstracts based on sentence level co-occurrence of KLK-7 and phenotype terms. Size of a circle indicates the square root of the number of publications covering a gene-phenotype pairing. Top 3 conditions with the highest specificity all of the skin-related diseases.

**Figure 5.**
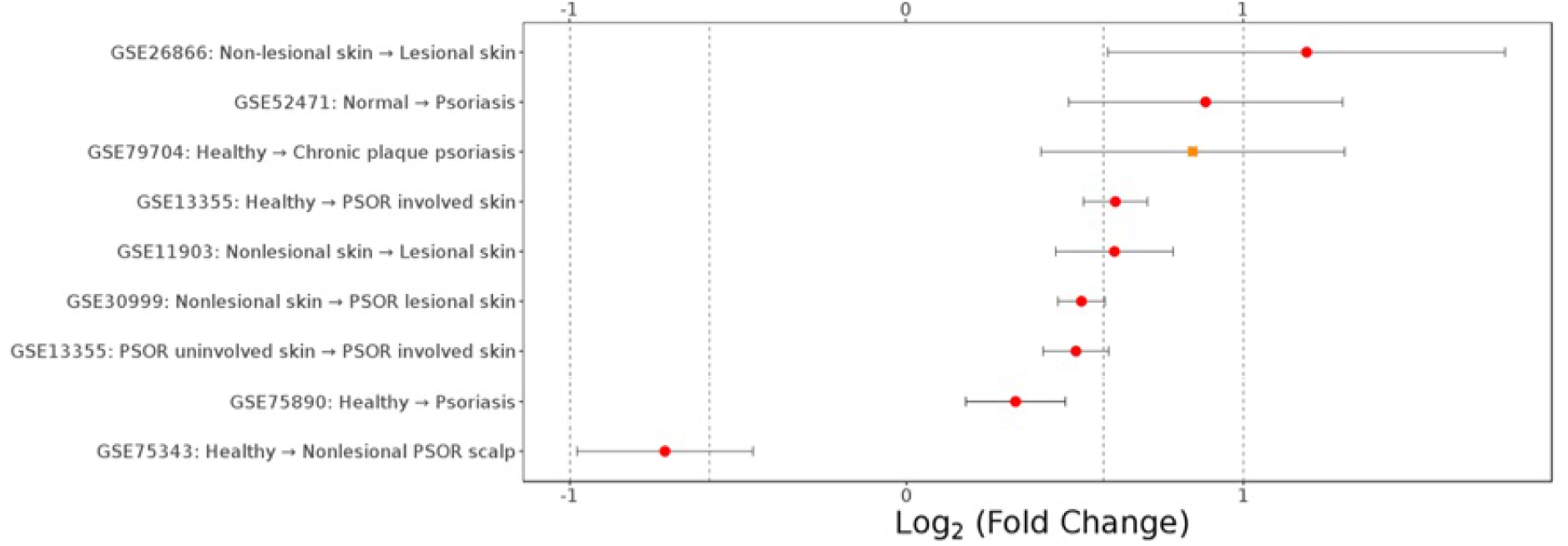
Survey of KLK-7 mRNA expression patterns in human psoriasis studies. For clarity, only statistically significant contrasts are shown (FDR<0.01). The fold change corresponds to the change from condition 1 to condition 2.

In the ankylosing spondylitis cohort, we discovered robust positive correlations of the disease-centric biomarkers LRG-1 and IL-6 with clinical severity. However, LRG-1 was upregulated upon apremilast treatment, providing molecular consistency to the clinical conclusion that the trial did not meet primary end points.

Although clear individual biomarker signals exist (IL-17A, KLK-7, IL-6, LRG-1), synchronized transitions by groups of analytes can capture the underlying biological complexity. Clinical trials present unique opportunities whereby these coordinated analytes are induced in pre-treated diseased conditions and subsequently followed in the presence of pharmacological treatment. There were two types of perturbation in our study design: longitudinal and pharmacological. Anchoring on both perturbation types, we were able to utilize time-series analyses to capture sub-group differences of apremilast pharmacodynamics. Analyte groups were catagorized by patterns to enable the identification of additional biomarkers with similar behavior. More specifically, we identified PEDF, MDC and ANGPTL4 from the same pharmacodynamic pattern as IL-17A and KLK-7, for which the combined rate of decline between Week 4 and 16 could differentiate apremilast clinical responses in psoriasis. To the best of our knowledge, this finding represents the only set of biomarkers with apremilast-treated clinical concordance thus far. We were unsuccessful in building post-hoc predictive models using baseline analytes for response status in any of the 3 trials, likely due to complex biology and small samples after sub-setting for timepoints/response groups (Supplementary Figure 4).

Another PDE4 inhibitor, roflumilast, was tested in vitro using sputum cells from patients with chronic obstructive pulmonary disease, but no effect on spontaneous MDC (CCL22) production was found^34^. There are no previous reports of apremilast or any other PDE4 inhibitor affecting PEDF, MDC and ANGPTL4 expression. Therefore, the findings here that apremilast can reduce levels of PEDF, MDC, and ANGPTL4 are novel and exemplify the learnings efforts from utilizing technology platforms in trials. We speculated that these subtle differences may shed light into shared biology between these related diseases.

Using 185,360 plasma biomarker measurements collected from 3 independent apremilast Phase III trials in ankylosing spondylitis, psoriasis and psoriatic arthritis, we found IL-17A and KLK-7 as robust biomarkers of psoriasis severity, which were also both biomarkers of general apremilast pharmacodynamic effects across 3 diseases. The steeper combined reduction rate of PEDF, KLK-7, MDC, ANGPTL4 from Week 4 to 16 differentiated responders from non-responders in the psoriasis trial. IL-6 and LRG-1 were identified as biomarkers with concordance to ankylosing spondylitis severity. Apremilast-induced LRG-1 increase was consistent with the conclusion that apremilast was ineffective in the ankylosing spondylitis trial. In addition to advancing the understandings of the apremilast therapeutic mechanism, these findings also help with the development of next-generation therapeutics. We demonstrated the value of, and advocate for the incorporation of exploratory molecular biomarker profiling into future clinical trials.

## Methods

### Response status definition

Subjects were considered to be responders at Week 16 if they met pre-defined criteria in each trial. In the ankylosing spondylitis trial, response was defined by a 20% improvement in the Assessment of SpondyloArthritis international Society (ASAS20) score. In the psoriasis trial, response was defined by a 75% improvement in Psoriasis Area and Severity Index (PASI) score (PASI75) or 50% improvement in PASI score (PASI50) and static Physician’s Global Assessment (sPGA) score of 0 or 1^35^. In the psoriatic arthritis trial, response was defined by a 20% improvement in the modified American College of Rheumatology (ACR20) response criteria^36^.

### Protein quantification

Protein profiling was performed by Myriad RBM that offered absolute protein quantitation in a CLIA certified laboratory with their MAP immunoassay panels for 150 analytes and Simoa ultrasensitive immunoassays for 5 analytes (Supplementary Table 9). In each trial, patient plasma samples were taken at each time point, and assayed with this platform. Pre-processing was performed on the raw data by removing individuals and proteins that had more than 50% missing values or values under the lower limit of quantitation (LLOQ). Remaining missing values were set to the protein average, and remaining <LLOQ values were set to 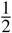 the protein LLOQ. Values were then log2 scaled and quantile normalized. In each trial, roughly 15% of the proteins were measured as <LLOQ in nearly all patients, and these proteins were largely consistent between the trials. This suggested possible mis-calibration of certain analytes to protein levels not observed in the data. Most proteins (80%) had no <LLOQ measurements, while about 5% had at least some <LLOQ measurements (between 0% and 50%). The final data contained 121 analytes in AS, 122 in PSOR and 155 in PSA trials.

### Correlation analyses with disease scores

Protein correlations from AS subjects were calculated for ASDAS, BASDAI, and BASFI. PASI total score was used for correlation in PSOR subjects, while DAS28 was used in the PSA trial. Lasso modelling was performed using *glmnet* package^37^ with 10-fold cross-validation. Univariate regression and multi-variate additive models were applied using R package *stats*, with age and gender as covariates. Associations were considered as significant with FDR less than 0.05 and *R*^2^ > 0.2. Aggregated expression change of IL-17A and KLK-7 between Week 4 and baseline were calculated using the corresponding beta coefficients: 

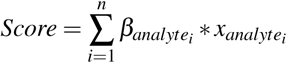

Difference in aggregated scores between groups were calculated by Mann-Whitney U test.

### Differential expression analysis

Differential expression analysis was performed using *limma* R package^38, 39^ with FDR < 0.05. Linear mixed-effect regression was performed when combining timepoints.

### Time-series analyses

Both CoGAPS and PatternMakers algorithms were applied using R package *CoGAPS*^29, 30, 39^. CoGAPS (Coordinated Gene Activity in Pattern Sets) is a Markov chain Monte Carlo Non-negative Matrix Factorization (NMF) method that quantifies distinct behavior patterns across different conditions (Figures 3A,C,E). Briefly, the method decomposes the input expression matrix into two matrices, pattern and the corresponding amplitude matrix, with minimal variance from the original matrix. The pattern matrix is the non-orthogonal basis represented by unit vectors. The vector coordinates of this basis represent the “relative strength” and can be interpreted as the differences in protein production comparable along the single vector. The magnitude of relative strength is a function of noise and reflects on relative protein expression levels to the mean of all analytes.

The input matrix for the current study was constructed as follows: mean expression values for each analyte were appended into a matrix, with analytes as rows, time points as columns, and separated by patient subgroups of comparisons. A similar standard error matrix was used to estimate goodness of fit by the CoGAPS module. Algorithm was run for 100 times with different random seeds, with minimum number of patterns equal to number of subgroups. To control stochastic nature of the algorithm we compared the mean *χ*^2^ between batch of runs with different number of iterations: 1000, 5000, 10000, 15000, 20000 (Supplemental Figure 6). Low value of *χ*^2^ represented precise matrix decomposition^40^. The number of iterations that resulted with the lowest mean *χ*^2^ was used for the subsequent analysis. Pairwise similarities between patterns of every CoGAPS run was calculated by root-mean-square-distance (RMSD). Patterns were merged if they met similarity criteria which was defined to be within 5% deviation of relative strength RMSD. Analyte associations to each pattern were calculated and ranked by the PatternMarkers algorithm. Analytes were assigned to a merged pattern if the occurrence frequency exceeded 50% across all of the runs.

### Systematic literature survey of KLK-7 and diseases

Sentence-level co-occurrence of KLK-7 and phenotypes terms from MEDLINE^®^abstracts (from January 1991 to June 2018) were extracted using SciBit^©^ v6.2 TERMite Expressions (TExpress) in Batch mode. A TExpress pattern specifying entity types including GENE and HPO was employed to ensure that synonyms of KLK-7 and all phenotype concepts from the Human Phenotype Ontology (HPO) were considered. Following extraction, a specificity score for each unique gene-phenotype co-occurrence was calculated based on mutual information between the gene and phenotype assessed by tf-idf scores, as described by Frijters et al.^41^ Specificity score reflected the relevancy of the association between the phenotype and the gene. Each unique phenotype was tagged with their highest HPO parent class(es) under phenotypic abnormality based on the HPO hierarchical relationships.

### Systematic mRNA survey of KLK-7 patterns in public psoriasis gene expression datasets

Relevant datasets were compiled, downloaded and re-analyzed from Gene Expression Omnibus (GEO)^42^. For each dataset, batch effects were explored and removed by the *SVA* package^39, 43^ *limma* package^38, 39^ was used for transcriptome-wide differential analyses.

## Data Availability

The data and scripts available at GitHub: https://github.com/imedvedeva-celgene/cogaps-apremilast. The ClinicalTrials.gov ID of the clinical studies used in the analysis were NCT01583374, NCT01172938, NCT01232283.

## Acknowledgements

The research was funded by Celgene Corporation. All analyte measurements were performed as in-kind contribution from Myriad RBM Inc. The authors are grateful to Dr. Xiaochun Ni for helpful discussions.

## Author contributions statement

D.E., S.T.L. M.W.B.T. and P.S. organized the study, I.V.M. and M.E.S. applied statistical analysis, J.A. contributed with systematic literature survey, R.Y. guided the research. I.V.M., P.S. and R.Y. wrote the manuscript. All authors read and approved the manuscript.

