## Supplemental Tables and Figures for "Large-scale Analyses of Disease Biomarkers and Apremilast Pharmacodynamic Effects" (from another version of this preprint: version 2, 2019-09-09)

Supplemental Material

Irina V. Medvedeva^1,*^, Matthew E. Stokes^1^, Dominic Eisinger^2^, Samuel T. LaBrie^2^, Jing Ai^1^, Matthew Trotter^3^, Peter Schafer^4^ and Robert Yang^1^

^1^Celgene Corporation, Informatics&Predictive Sciences, Cambridge, 02140, USA

^2^Myriad RBM Inc., Austin, 78759, USA

^3^Celgene Corporation, Celgene Institute for Translational Research Europe (CITRE), Sevilla, 41092, Spain

^4^Celgene Corporation, Translational Development, Summit, 07901, USA

^*^

**Tables**

| *Table 1. List of significant coefficients from univariate regression model built for changes in analyte’s production between Week 4 and Week 0 and PASI total score for PSOR patients* | | | | | | | |
| --- | --- | --- | --- | --- | --- | --- | --- |
|  | **Beta (analytes)** | **p-value (analytes)** | **Beta (gender)** | **p-value (gender)** | **Beta (age)** | **p-value (age)** | **p-value (F)** |
| **E-Selectin** | 5.125328 | 1.46E-02 | 2.671034 | 0.07354241 | 0.04565763 | 0.2929972 | 1.61E-02 |
| **MDC** | 2.567925 | 1.46E-02 | 2.320716 | 0.07354241 | 0.03253138 | 0.3738943 | 1.61E-02 |
| *Table 2. List of significant coefficients from univariate regression model built for changes in analyte’s production between Week 16 and Week 4 and PASI total score for PSOR patients* | | | | | | | |
|  | **Beta (analytes)** | **p-value (analytes)** | **Beta (gender)** | **p-value (gender)** | **Beta (age)** | **p-value (age)** | **p-value (F)** |
| **IL-17A** | 3.003621 | 1.02E-02 | 1.0312 | 0.4610179 | -0.0127957 | 0.7861655 | 7.42E-02 |
| **KLK-7** | 4.12521 | 3.43E-03 | 0.6719603 | 0.5676315 | -0.0102978 | 0.8182355 | 2.93E-02 |
| **MDC** | 2.998181 | 2.98E-02 | 0.7127427 | 0.564288 | -0.0354486 | 0.5567633 | 1.83E-01 |
| *Table 3. List of significant coefficients from univariate regression model built for changes in analyte’s production between Week 16 and Week 0 and PASI total score for PSOR patients* | | | | | | | |
|  | **Beta (analytes)** | **p-value (analytes)** | **Beta (gender)** | **p-value (gender)** | **Beta (age)** | **p-value (age)** | **p-value (F)** |
| **MDC** | 4.594724 | 7.73E-06 | 3.079595 | 0.0639386 | -0.0038608 | 0.9379449 | 2.65E-05 |
| **IL-17A** | 3.812114 | 3.31E-05 | 1.542033 | 0.24794 | 0.07616996 | 0.8140634 | 1.02E-04 |
| **KLK-7** | 4.35861 | 5.30E-05 | 2.893446 | 0.0639386 | 0.04064567 | 0.8140634 | 1.55E-04 |
| **E-Selectin** | 7.126629 | 9.55E-04 | 3.695473 | 0.0639386 | 0.01702296 | 0.8140634 | 2.23E-03 |
| **IL-16** | 3.344772 | 3.09E-03 | 3.180763 | 0.0639386 | 0.02723837 | 0.8140634 | 6.39E-03 |
| **TN-C** | 4.119571 | 1.89E-02 | 2.93368 | 0.0639386 | 0.06098382 | 0.8140634 | 3.18E-02 |
| **Progranulin** | 3.53925 | 4.73E-02 | 3.723086 | 0.0639386 | 0.01462927 | 0.818219 | 6.95E-02 |

| *Table 4. List of significant coefficients from univariate regression model built for changes in analyte’s production between Week 4 and Week 0 and ASDAS score for AS patients* | | | | | | | |
| --- | --- | --- | --- | --- | --- | --- | --- |
|  | **Beta (analytes)** | **p-value (analytes)** | **Beta (gender)** | **p-value (gender)** | **Beta (age)** | **p-value (age)** | **p-value (F)** |
| **LRG1** | 0.5048268 | 6.97E-09 | -0.10979315 | 0.2851051 | -0.02633521 | 0.5202252 | 1.17E-07 |
| **AGP-1** | 0.2508537 | 1.09E-02 | -0.07069833 | 0.4081878 | -0.03177247 | 0.5202252 | 6.81E-02 |
| **IL-18bp** | 0.2728842 | 1.09E-02 | -0.08925593 | 0.3069795 | -0.02649034 | 0.5249954 | 6.81E-02 |
| **IL-6** | 0.2244431 | 4.71E-02 | -0.1055207 | 0.2851051 | -0.03908487 | 0.5202252 | 2.28E-01 |
| **Prostasin** | -0.214788 | 4.71E-02 | -0.10144949 | 0.2851051 | -0.02632393 | 0.5249954 | 2.28E-01 |
| *Table 5. List of significant coefficients from univariate regression model built for changes in analyte’s production between Week 16 and Week 4 and ASDAS score for AS patients* | | | | | | | |
|  | **Beta (analytes)** | **p-value (analytes)** | **Beta (gender)** | **p-value (gender)** | **Beta (age)** | **p-value (age)** | **p-value (F)** |
| **LRG1** | 0.6350613 | 1.07E-10 | -0.00102202 | 0.9995394 | 0.05406444 | 0.2381342 | 1.95E-09 |
| **AGP-1** | 0.4532631 | 3.68E-08 | -0.06725939 | 0.9995394 | 0.05899935 | 0.2214485 | 5.25E-07 |
| **IL-6** | 0.4101409 | 2.06E-05 | 0.032781562 | 0.9995394 | 0.05606142 | 0.2448171 | 2.24E-04 |
| **TNF RI** | 0.4374841 | 1.46E-03 | -0.00651405 | 0.9995394 | 0.08996539 | 0.2132886 | 1.03E-02 |
| **BAFF** | 0.1861229 | 4.66E-02 | -0.0343849 | 0.9995394 | 0.07288844 | 0.2132886 | 1.99E-01 |
| **Neurop-1** | 0.2056688 | 4.66E-02 | -0.0061204 | 0.9995394 | 0.05971983 | 0.2377463 | 1.99E-01 |
| **PEDF** | 0.2992843 | 4.66E-02 | 0.001364536 | 0.9995394 | 0.07838157 | 0.2132886 | 1.99E-01 |
| **TATI** | 0.29163 | 4.66E-02 | -0.00712883 | 0.9995394 | 0.06804483 | 0.2132886 | 1.99E-01 |
| *Table 6. List of significant coefficients from univariate regression model built for changes in analyte’s production between Week 16 and Week 0 and ASDAS score for AS patients* | | | | | | | |
|  | **Beta (analytes)** | **p-value (analytes)** | **Beta (gender)** | **p-value (gender)** | **Beta (age)** | **p-value (age)** | **p-value (F)** |
| **LRG1** | 0.5571898 | 3.84E-08 | -0.12209829 | 0.3518272 | 0.030388863 | 0.5746815 | 5.58E-07 |
| **AGP-1** | 0.3259233 | 2.43E-04 | -0.10748969 | 0.3518272 | 0.024357368 | 0.6450627 | 2.09E-03 |
| **IL-6S** | 0.3050261 | 2.99E-03 | -0.10551392 | 0.3518272 | 0.008643994 | 0.8687516 | 2.00E-02 |
| **MPIF-1** | 0.268508 | 8.84E-03 | -0.08885186 | 0.4166016 | 0.042679277 | 0.5504894 | 4.86E-02 |
| **SCFR** | -0.302818 | 8.84E-03 | -0.13443356 | 0.3518272 | 0.023398262 | 0.6598157 | 4.86E-02 |
| **TNF RI** | 0.361627 | 8.84E-03 | -0.1196068 | 0.3518272 | 0.047785724 | 0.5504894 | 4.86E-02 |
| **RBP-4** | -0.2698329 | 1.60E-02 | -0.09714638 | 0.381725 | 0.030123356 | 0.5892207 | 7.99E-02 |
| **SAA** | 0.173719 | 2.62E-02 | -0.11518927 | 0.3518272 | 0.03140821 | 0.5779453 | 1.20E-01 |
| **HE4** | -0.2611998 | 4.35E-02 | -0.14521165 | 0.3518272 | 0.0436594 | 0.5504894 | 1.83E-01 |

*Table 7. ANOVA comparison of the nested linear models predicting PASI score including listed proteins and age and gender as covariates for PSOR patients at baseline. P-values represents the model comparison between additive model and each of the univariate models.*

| **model** | **R^2^** | **RSS** | **p** |
| --- | --- | --- | --- |
| *Univariate IL-17A* | 0.29 | 3558 | 5.10E-04 |
| *Univariate KLK-7* | 0.26 | 3706 | 3.97E-05 |
| *Additive model: IL-17A+KLK-7* | 0.35 | 3210 | NA |

*Table 8. ANOVA comparison of the nested linear models predicting ASDAS score including listed proteins and age and gender as covariates for AS patients at baseline. P-values represents the model comparison between additive model and each of the univariate models.*

| **model** | **R^2^** | **RSS** | **p** |
| --- | --- | --- | --- |
| *Univariate IL-6* | 0.30 | 112 | 3.59E-18 |
| *Univariate LRG1* | 0.46 | 86 | 2.74E-04 |
| *Additive model: IL-6+LRG1* | 0.49 | 83 | NA |

*Table 9. Plasma Proteomic Biomarkers Tested*

| **Multiplex Bead-Based MAP Immunoassays** | | |
| --- | --- | --- |
| 6Ckine | Gelsolin | Neuronal Cell Adhesion Molecule (Nr-CAM) |
| Alpha-1-acid glycoprotein 1 (AGP-1) | Granulocyte Colony-Stimulating Factor (G-CSF) | Neuropilin-1 |
| Alpha-1-Antitrypsin (AAT) | Growth/differentiation factor 15 (GDF-15) | Neutrophil Activating Peptide 2 (NAP-2) |
| Alpha-1-Microglobulin (A1Micro) | Growth-Regulated alpha protein (GRO-alpha) | Osteocalcin |
| Amphiregulin (AR) | Haptoglobin | Osteoprotegerin (OPG) |
| Angiogenin | HE4 | P-Selectin |
| Angiopoietin-1 (ANG-1) | Hemopexin | Pancreatic secretory trypsin inhibitor (TATI) |
| Angiopoietin-2 (ANG-2) | Heparin-Binding EGF-Like Growth Factor (HB-EGF) | Pepsinogen I (PGI) |
| Angiopoietin-related protein 4 (ANGPTL4) | Hepsin | Periostin |
| Antileukoproteinase (ALP) | Immunoglobulin A (IgA) | Pigment Epithelium Derived Factor (PEDF) |
| Antithrombin-III (AT-III) | Immunoglobulin E (IgE) | Placenta Growth Factor (PLGF) |
| Apolipoprotein(a) (Lp(a)) | Immunoglobulin M (IgM) | Platelet endothelial cell adhesion molecule (PECAM-1) |
| AXL Receptor Tyrosine Kinase (AXL) | Insulin-like Growth Factor-Binding Protein 2 (IGFBP-2) | Platelet-Derived Growth Factor BB (PDGF-BB) |
| B cell-activating factor (BAFF) | Insulin-like Growth Factor-Binding Protein 7 (IGFBP-7) | Progranulin |
| Beta-2-Microglobulin (B2M) | Interferon alpha (IFN-alpha) | Prostasin |
| Betacellulin (BTC) | Interferon gamma Induced Protein 10 (IP-10) | Prostate-Specific Antigen, Free (PSA-f) |
| C-Reactive Protein (CRP) | Interferon-inducible T-cell alpha chemoattractant (ITAC) | Pulmonary surfactant-associated protein D (SP-D) |
| Cadherin-1 (E-Cad) | Interleukin-1 receptor type 1 (IL-1RI) | Receptor for advanced glycosylation end products (RAGE) |
| Cancer Antigen 15-3 (CA-15-3) | Interleukin-1 receptor type 2 (IL-1RII) | Retinol-binding protein 4 (RBP-4) |
| Carbonic anhydrase 9 (CA-9) | Interleukin-2 receptor alpha (IL-2 receptor alpha) | S100 calcium-binding protein B (S100-B) |
| Carcinoembryonic antigen-related cell adhesion molecule 1 (CEACAM1) | Interleukin-6 receptor (IL-6r) | Sclerostin |
| Cathepsin D | Interleukin-6 receptor subunit beta (IL-6R beta) | Serum Amyloid A Protein (SAA) |
| CD27 antigen (CD27) | Interleukin-13 (IL-13) | Sex Hormone-Binding Globulin (SHBG) |
| CD40 Ligand (CD40-L) | Interleukin-16 (IL-16) | Sortilin |
| CD163 | Interleukin-18-binding protein (IL-18bp) | ST2 |
| Chemokine CC-4 (HCC-4) | Interleukin-22 (IL-22) | Stromal cell-derived factor-1 (SDF-1) |
| Chromogranin-A (CgA) | Interleukin-31 (IL-31) | Superoxide Dismutase 1, soluble (SOD-1) |
| Ciliary Neurotrophic Factor (CNTF) | Kallikrein-7 (KLK-7) | T Lymphocyte-Secreted Protein I-309 (I-309) |
| Clusterin (CLU) | Lactoferrin (LTF) | Tenascin-C (TN-C) |
| Complement C3 (C3) | Latency-Associated Peptide of Transforming Growth Factor beta 1 (LAP TGF-b1) | Tetranectin |
| Cystatin-B | Leucine-rich alpha-2-glycoprotein (LRG1) | Thrombin-Activatable Fibrinolysis (TAFI) |
| Cystatin-C | Macrophage-Derived Chemokine (MDC) | Thrombospondin-1 |
| Decorin | Macrophage Inflammatory Protein-3 alpha (MIP-3 alpha) | Thymus and activation-regulated chemokine (TARC) |
| Dickkopf-related protein 1 (DKK-1) | Macrophage inflammatory protein 3 beta (MIP-3 beta) | Thymus-Expressed Chemokine (TECK) |
| E-Selectin | Macrophage Migration Inhibitory Factor (MIF) | Thyroxine-Binding Globulin (TBG) |
| Eotaxin-2 | Macrophage-Stimulating Protein (MSP) | Tissue Inhibitor of Metalloproteinases 2 (TIMP-2) |
| Eotaxin-3 | Maspin | Tissue Inhibitor of Metalloproteinases 3 (TIMP-3) |
| Epidermal Growth Factor (EGF) | Mast/stem cell growth factor receptor (SCFR) | TNF-Related Apoptosis-Inducing Ligand Receptor 3 (TRAIL-R3) |
| Epidermal Growth Factor Receptor (EGFR) | Matrix Metalloproteinase-1 (MMP-1) | Transforming Growth Factor beta-3 (TGF-beta-3) |
| Epiregulin (EPR) | Matrix Metalloproteinase-2 (MMP-2) | Tumor necrosis factor ligand superfamily member 12 (Tweak) |
| Epithelial-Derived Neutrophil-Activating Protein 78 (ENA-78) | Matrix Metalloproteinase-7 (MMP-7) | Tumor necrosis factor ligand superfamily member 13 (APRIL) |
| Erythropoietin (EPO) | Matrix Metalloproteinase-9, total (MMP-9, total) | Tumor Necrosis Factor Receptor I (TNF RI) |
| Fas Ligand (FasL) | Matrix Metalloproteinase-10 (MMP-10) | Urokinase-type plasminogen activator receptor (uPAR) |
| FASLG Receptor (FAS) | Mesothelin (MSLN) | Vascular endothelial growth factor D (VEGF-D) |
| Fatty Acid-Binding Protein, liver (FABP, liver) | Monocyte Chemotactic Protein 2 (MCP-2) | Vascular Endothelial Growth Factor Receptor 2 (VEGFR-2) |
| Fibrinogen | Monocyte Chemotactic Protein 3 (MCP-3) | Vitamin D-Binding Protein (VDBP) |
| Fibroblast Growth Factor 21 (FGF-21) | Monocyte Chemotactic Protein 4 (MCP-4) | Vitronectin |
| Fibroblast growth factor 23 (FGF-23) | Myeloid Progenitor Inhibitory Factor 1 (MPIF-1) | von Willebrand Factor (vWF) |
| Fibulin-1C (Fib-1C) | Myeloperoxidase (MPO) | Bone morphogenetic protein 9 (BMP-9) |
| Ficolin-3 | Nerve Growth Factor beta (NGF-beta) | YKL-40 |
| **Simoa Bead-Based Ultrasensitive Immunoassays** | | |
| B Lymphocyte Chemoattractant Simoa (BLC Simoa) | Interleukin-6 Simoa (IL-6 Simoa) | Tumor Necrosis Factor alpha Simoa (TNFa Simoa) |
| Interleukin-12 Subunit p40 Simoa (IL-12p40 Simoa) | Interleukin-17A Simoa (IL-17A Simoa) |  |

**Figures**

Week 16

Week 4

Baseline

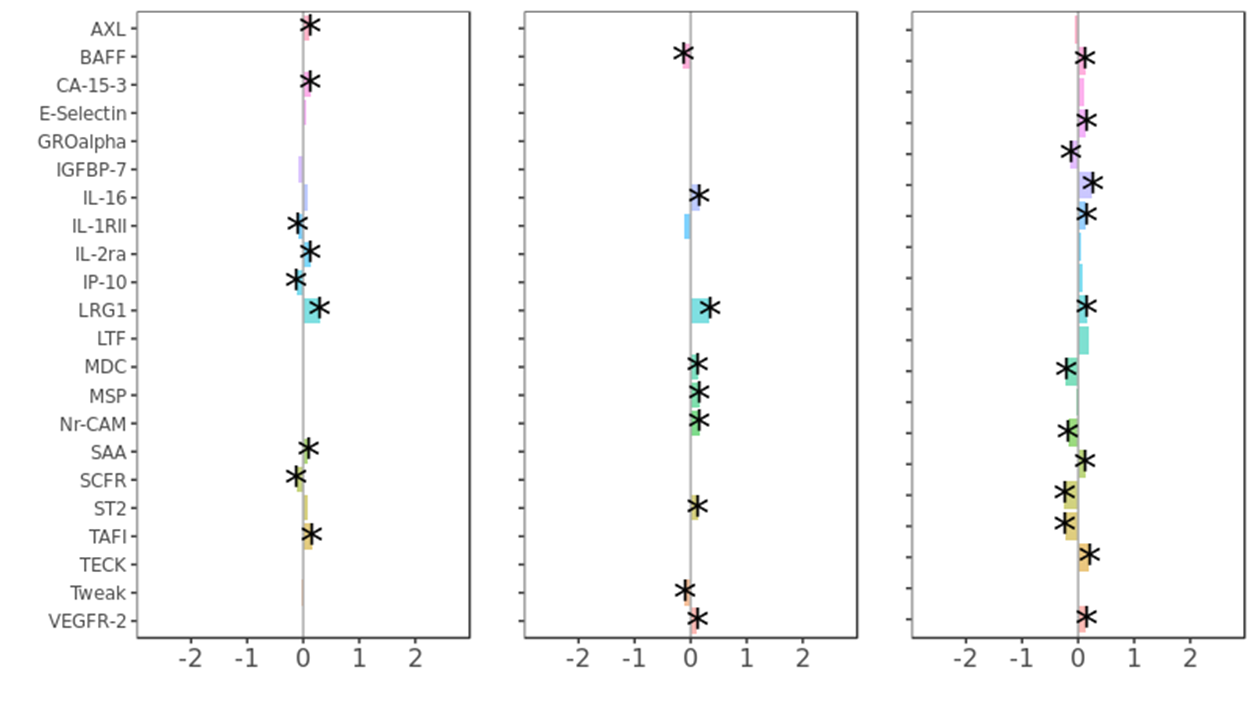

Figure 1. The value of beta coefficients in lasso modeling of protein production estimates predicting ASDAS score in patients with ankylosing spondylitis in three time points. Stars indicates the proteins that were significant in the model.

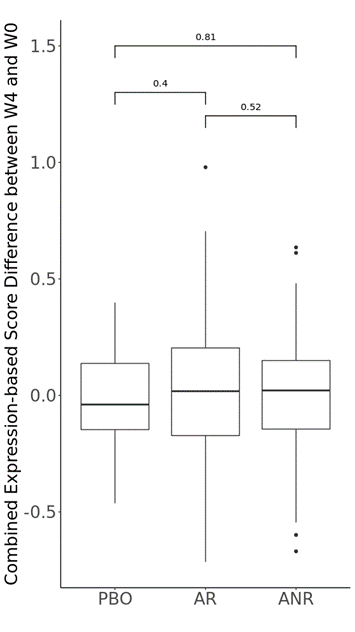

*Figure 2. Mann-Whitney comparison of combined expression-based score difference of IL-6 and LRG1 at Week 0 and Week 4 in different groups: placebo (PBO), apremilast responders (AR) and apremilast non-responders (ANR). P-values are shown above, no significance was found.*

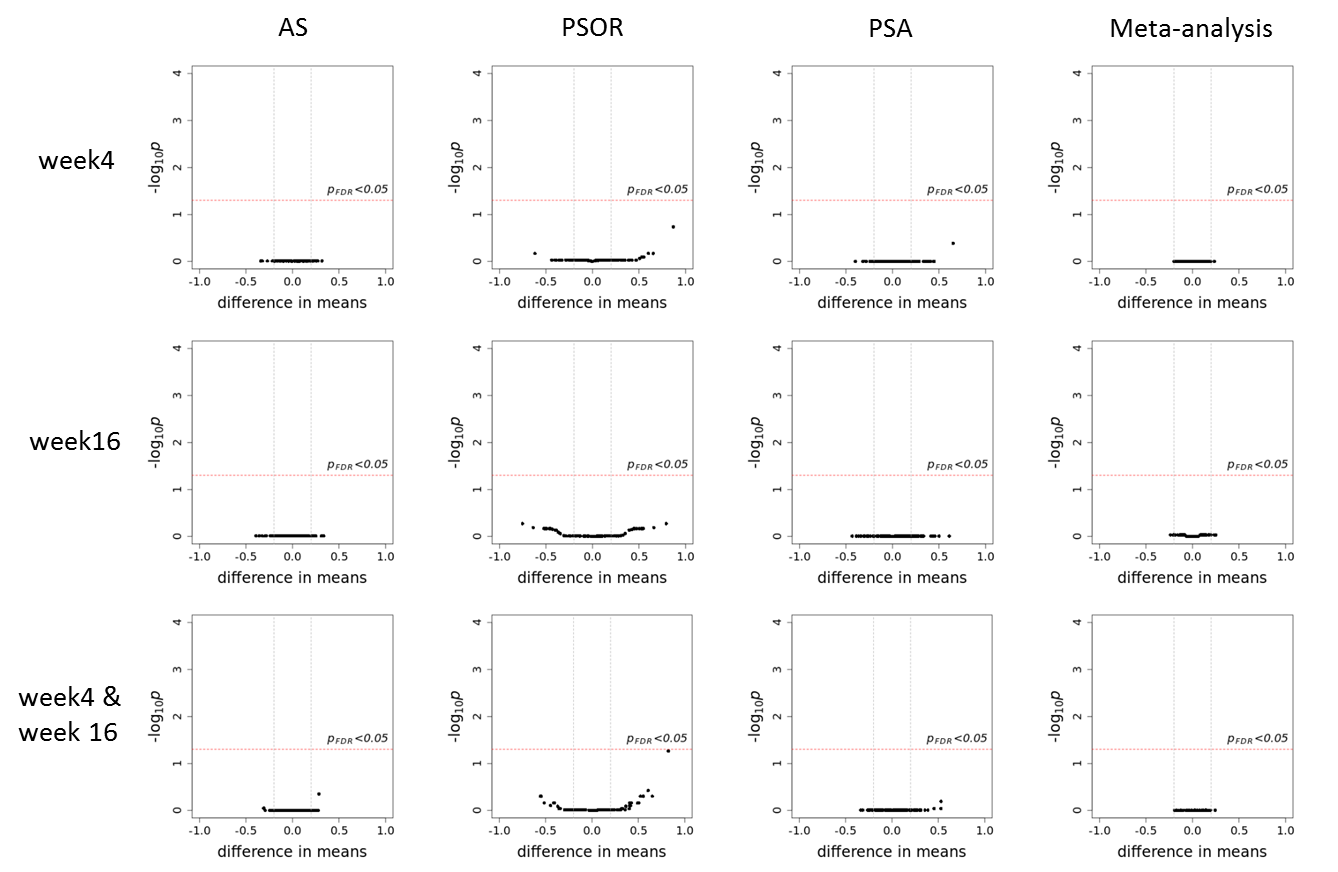

Figure 3. Differential expression analysis between apremilast non-responders and apremilast responders arms for specified time points and meta-analysis across all diseases. The threshold corresponding to the p-value 0.05 is shown as red line. None of the proteins reached the significance.

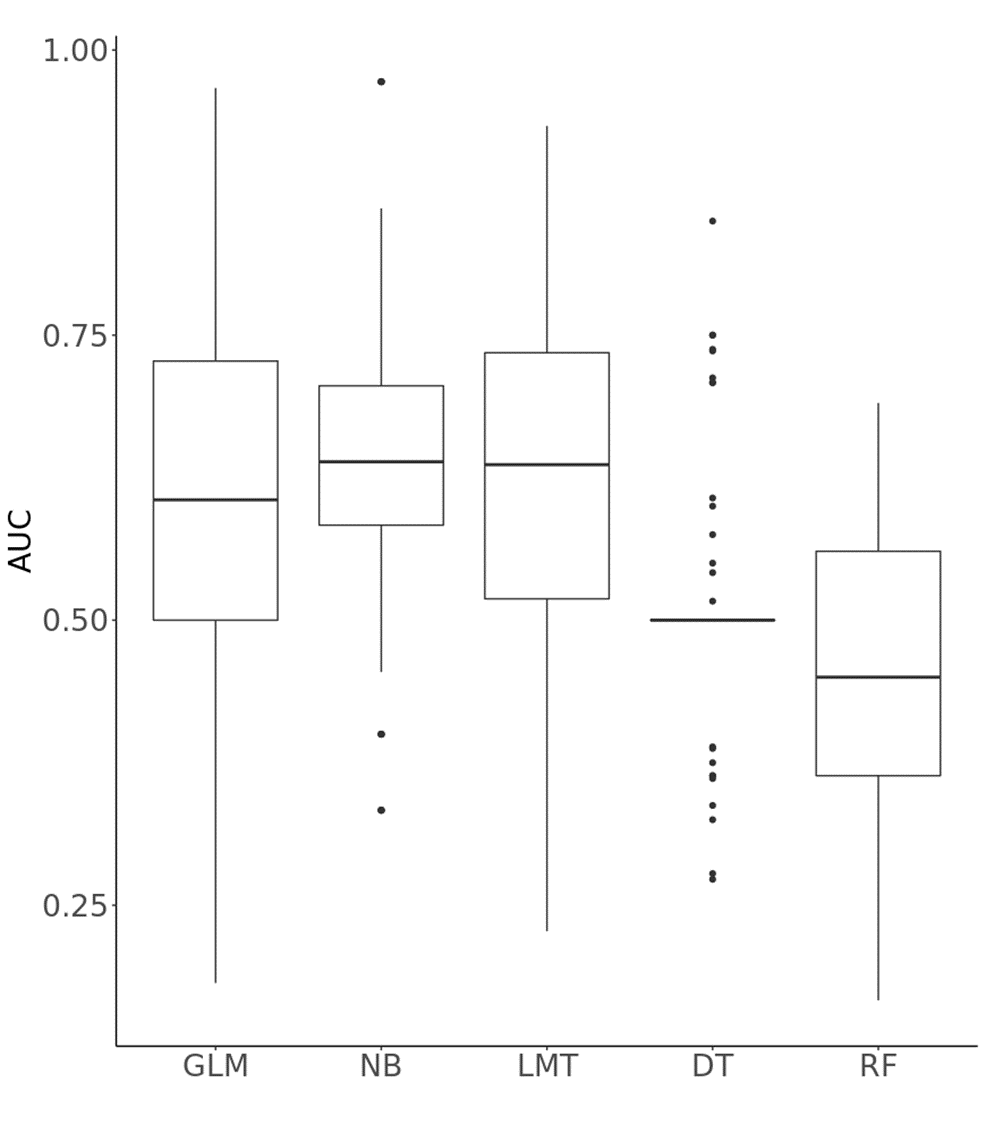

*Figure 4. Prediction models using KLK7, PEDF, MDC, ANGPTL4 analytes associated with downregulative pattern in CoGAPS analysis in psoriasis responders. AUC was used to measure the effectiveness of prediction on 80% of dataset after 100 random split of the samples. Models used: GLM – global linear model, NB – naïve Bayes, LMT – logistic model tree, DT – decision tree, RF – random forest.*

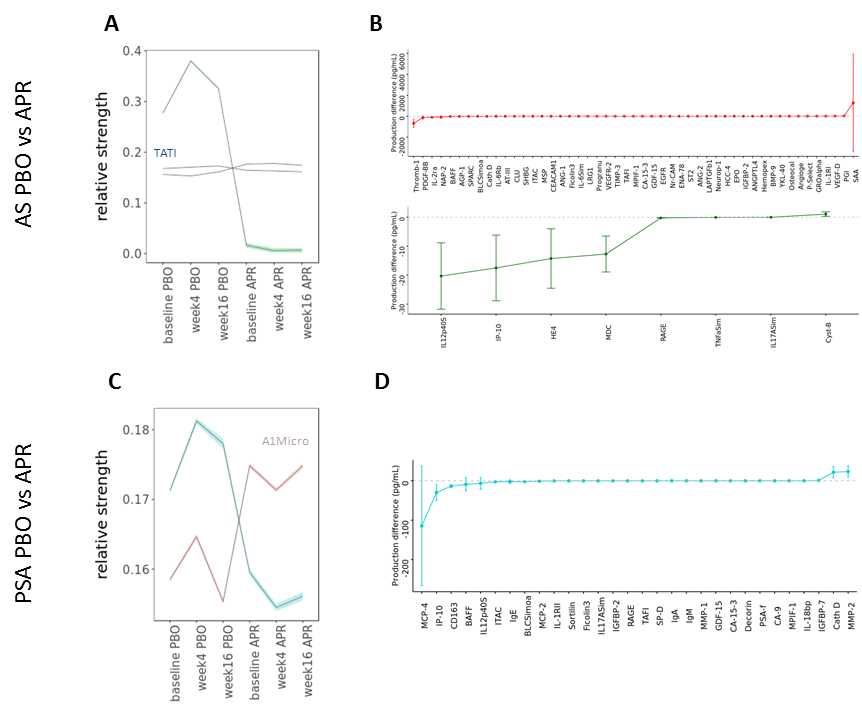

Figure 5. CoGaps patterns (A,C) and corresponding proteins assigned to each pattern (B, D) and ranked according to mean difference of production between week 4 and baseline. Colors of plots in (B,D) match the colors of the patterns. The patterns represented by single proteins are shown: blue pattern with TATI in AS, red pattern with A1Micro in PSA. Green pattern in AS and blue pattern in PSA represent the downregulation under apremilast treatment. AS - ankylosing spondylitis; PSA - psoriatic arthritis; PBO - placebo treatment arm; APR - apremilast treatment arm.

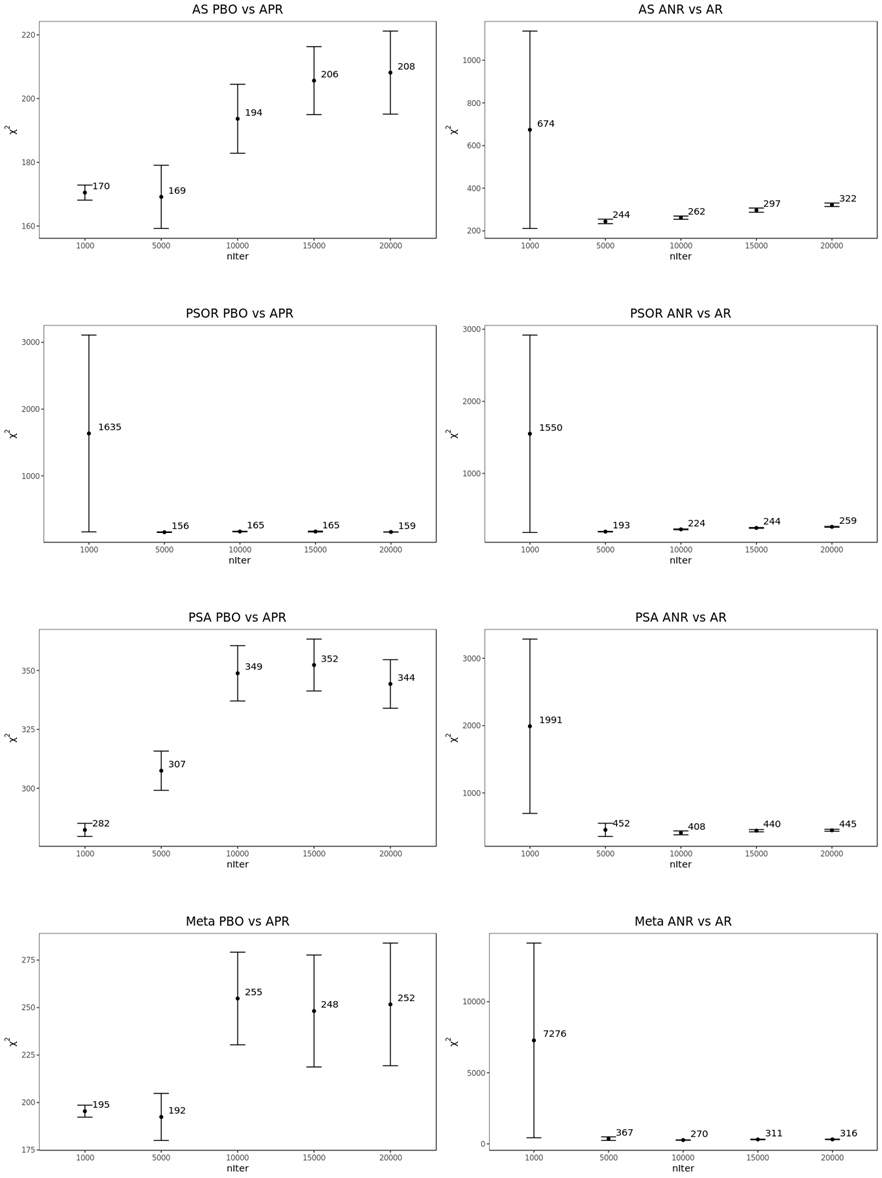

Figure 6. Mean χ^2^-square value for the different number of iterations in CoGAPS for different condition tests. AS - ankylosing spondylitis; PSOR - psoriasis; PSA - psoriatic arthritis; PBO - placebo treatment arm; APR - apremilast treatment arm; ANR – apremilast non-responders; AR – apremilast responders; Meta – meta-analysis of AS, PSOR and PSA trials combined.
